# Short-term exposure to culture media used in human ART shapes early calcium oscillations in ICSI-fertilized mouse oocytes and impacts adult phenotype

**DOI:** 10.1101/2025.03.14.643329

**Authors:** Bernadette Banrezes, Thierry Sainte Beuve, Anne Frambourg, Alice Jouneau

## Abstract

**Purpose:** The influence of culture media used during in vitro fertilization (IVF) on offspring phenotype remains controversial. However, specific effects of short exposure time after fertilization remain underexplored. By evaluating Ca^2+^ oscillations as a readout of the first response of eggs to their microenvironment, we aim to investigate if early differences correlate with later adult phenotypes.

**Methods:** Oocytes fertilized by ICSI were cultured for four hours in three different media (Cook and Vitrolife, used in human IVF, and KSOM, used for mouse embryos). They were either measured for Ca^2+^ oscillations or transferred into pseudo-pregnant females. After birth, growth curves of pups were measured up to adulthood and various organs weighed.

**Results:** Culture media significantly modulate Ca^2+^ oscillations during oocyte activation. ICSI-fertilized oocytes cultured in Cook and Vitrolife exhibited fewer oscillations, lower frequency, and reduced variability compared to KSOM. These early differences correlated with long-term developmental outcomes: females from Cook and Vitrolife cultures were heavier throughout growth and had larger adult organ sizes compared to those from KSOM.

**Conclusions:** Brief exposure to media immediately after ICSI shapes Ca^2+^ dynamics and adult phenotypes. Optimizing embryo culture protocols in assisted reproductive technologies may improve IVF outcomes by modulating metabolic pathways linked to development.

## 1. INTRODUCTION

The number of assisted reproductive technology (ART) treatment cycles continues to increase, reflecting growing reliance on in vitro fertilization (IVF) and related techniques^1,2^. Among IVF techniques, intracytoplasmic sperm injection (ICSI) has become the dominant method, accounting for 72.7% of procedures in Europe^1^ and 75.7% in the United States^3^. Given the widespread use of ICSI, it is essential to understand how the different culture media used immediately after fertilization can influence the development of ICSI-fertilized oocytes.

ICSI involves the direct injection of a single sperm into an oocyte, triggering a cascade of Ca^2+^ oscillations that activate mitochondrial metabolism and stimulate embryo development^4,5^. As mitochondrial function is critical during early embryogenesis, the surrounding culture environment may play a key role in modulating these metabolic processes. The composition of the embryo culture medium, as well as the duration of exposure, may influence implantation potential and long-term developmental outcomes^6,7^

Despite the recognized importance of embryo culture conditions, their precise role in shaping long-term development remains debated. Several human studies suggest that differences in the composition of embryonic culture media are associated with variations in birth weight, two-year weight, and up to nine years of age in children conceived by ART^8–11^. However, other studies report no significant association^12–14^. These conflicting findings suggest that additional factors— such as differences in medium formulation, fertilization technique, and patient variability—may contribute to outcome variability, underscoring the need for standardized ART laboratory protocols^15^. In this context, understanding how early Ca^2+^ oscillation patterns relate to long-term phenotypes is clinically relevant, as it may help explain phenotypic variability among ART-conceived individuals and guide improvements in embryo culture protocols.

In animal models, the composition of culture media has been shown to influence metabolic activity, modulate developmental trajectories, and even shape adult body weight^16–21^. However, one of the main challenges in evaluating the functional impact of commercial embryo culture media is the lack of transparency regarding their exact chemical composition. Relevant studies have identified significant differences in the concentrations of more than 39 chemical components across sixteen commercial human embryo culture media^22,23^. A more recent analysis of fifty-six human embryo culture media^24^ has further raised concerns about how this variability may influence embryonic responses and long-term developmental potential^8^.

Before being marketed, embryo culture media undergo safety testing, such as the Mouse Embryo Assay (MEA), which ensures that at least 80% of mouse embryos fertilized in vivo reach the blastocyst stage^25^. However, the MEA has notable limitations: it does not assess ICSI-fertilized oocytes or examine the long-term effects of culture media on embryonic development. Given the increasing reliance on ICSI, it is crucial to evaluate how post-fertilization culture conditions influence both early embryonic response and long-term phenotypic outcomes.

A key determinant during this critical post-fertilization period is the regulation of mitochondrial activation and genome reprogramming by Ca^2+^ oscillations^4,26–28^. These oscillations, which persist for approximately four hours before ceasing at pronucleus (PN) formation, serve as a real-time indicator of how embryos respond to their culture media formulation. These findings emphasize the need to integrate alternative assessment methods into the MEA, incorporating both early metabolic markers and long-term phenotypic evaluations to improve ART outcome prediction.

To address these knowledge gaps, we investigated how different ART culture media affect the initial metabolic responses of mouse oocytes after ICSI and their long-term developmental effects. Specifically, we compared two widely used ART media - Cook and Vitrolife - one of which has been associated in humans with differences in birth weight, weight at two years old, and even up to nine years of age in children conceived by ART^8–11^. As a reference, we used KSOM, a well-established mouse embryo culture medium^29^.

We first analyzed Ca^2+^ oscillation patterns in ICSI-fertilized oocytes cultured for four hours in these three culture media. We then evaluated whether these early metabolic responses correlated with in vivo developmental potential following embryo transfer. Key parameters included Ca^2+^ oscillation dynamics, embryonic development rates, postnatal growth and organ and tissue weights.

By integrating these analyses, we aimed to determine whether early Ca^2+^ responses in ICSI-fertilized oocytes could serve as predictive markers of long-term developmental potential. This study provides new insights into how human embryo culture media composition influences a critical four-hour post-fertilization window and its lasting effects on offspring phenotype.

## 2. MATERIALS AND METHODS

### 2.1. Gametes production and collection

Mature oocytes were recovered from superovulated B6CBAF1/JRj female mice (Janvier Labs, France) as described previously in Ozil *et al.*^18^. Metaphase II oocytes were collected in a Petri dish (Greiner Bio-One) containing hyaluronidase (90 i.u. ml^-1^, H3506, Sigma) in Hepes-buffered KSOM medium^29^ to remove cumulus cells. After washing and selection, the oocytes were placed in a drop of KSOM medium under mineral oil (M8410, Sigma) and incubated in a 5% CO_2_ atmosphere at 37°C. Spermatozoa from B6CBAF1/JRj males (Janvier Labs, France) were extracted from the caudal epididymis on each day of the experiment and suspended in KSOM medium^18^. This study was approved by the French Ministry of Higher Education, Research, and Innovation and by the local Ethics Committee for Animal Experiments at the author’s institution, under approval numbers no.12-165 and no.16-22 INRAE. All animal procedures were performed in accordance with the Code for Methods and Welfare Considerations in Behavioral Research (Directive 2010/63/EU).

### 2.2. Culture media formulation

The *in vitro* culture was carried out either in three different media. Cook and Vitrolife media were obtained from their respective suppliers, Cook Medical (K-SICM, Cook group, Ireland) and Vitrolife (G1™ Version 5 PLUS, Vitrolife group, Sweden). The compositions of these two commercial media were analysed for concentrations by Morbeck *et al.*^22^ and presented in Tables 1&2. The KSOM was made in the laboratory^29^. All media contained proteins, human serum albumin (HSA) already mixed in the Cook and Vitrolife, and bovine serum albumin (BSA, A3311, Sigma) added to the KSOM (see Table 1). The KSOM contains no amino acids (see Table 2). All inorganic and organic components were sourced from Sigma Chemical Company.

**Table 1:**
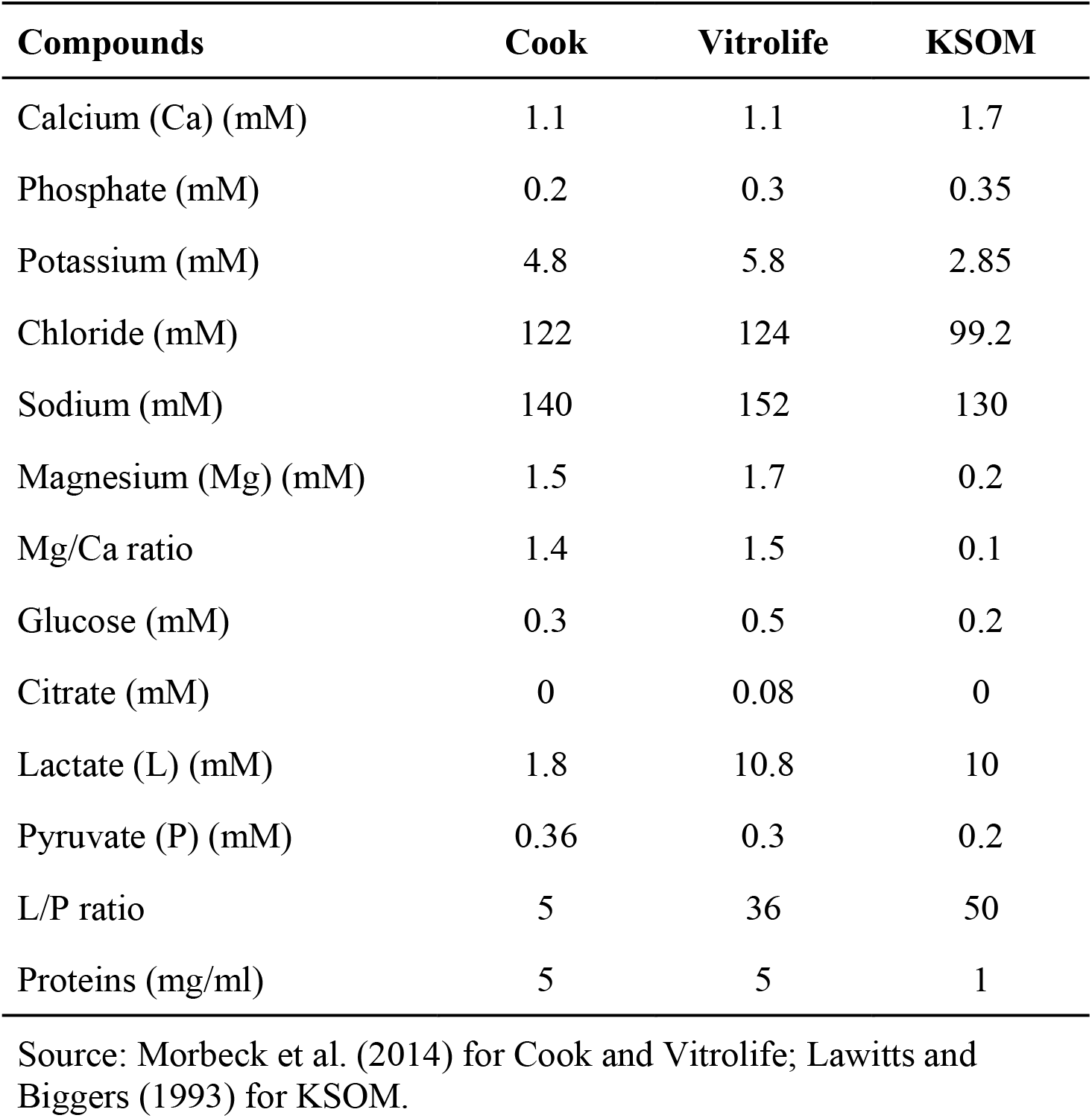
Comparison of electrolytes, carbohydrates and proteins in embryo culture media.

**Table 2:** Comparison of amino acids in embryo culture media.

| Compounds | Cook | Vitrolife | KSOM |
| --- | --- | --- | --- |
| Essential (μM) |  |  |  |
| Arg | 25 | 0 | 0 |
| Cys | 2 | 0 | 0 |
| His | 8 | 0 | 0 |
| Ile | 17 | 0 | 0 |
| Leu | 18 | 0 | 0 |
| Lys | 18 | 0 | 0 |
| Met | 4 | 0 | 0 |
| Phe | 8 | 0 | 0 |
| Thr | 18 | 0 | 0 |
| Trp | 2 | 0 | 0 |
| Tyr | 12 | 0 | 0 |
| Val | 17 | 0 | 0 |
| Nonessential (μM) |  |  |  |
| Ala | 135 | 148 | 0 |
| Asn | 88 | 126 | 0 |
| Asp | 81 | 0 | 0 |
| Glu | 90 | 0 | 0 |
| Gln | 30 | 0 | 0 |
| Gly | 6647 | 135 | 0 |
| Pro | 85 | 112 | 0 |
| Ser | 92 | 127 | 0 |
| Tau | 6489 | 131 | 0 |

Source: Morbeck et al. (2014) for Cook and Vitrolife; Lawitts and Biggers (1993) for KSOM.

### 2.3. Intra-cytoplasmic sperm injection

Oocytes were fertilized via ICSI at room temperature using a NIKON TMD microscope (NIKON, Japan), following the protocol described by Ozil et al.^18^. After sperm head injection, the plasma membrane of the oocyte was mechanically compressed against the zona pellucida to ensure sealing and prevent ion influx during membrane repair. ICSI-fertilized oocytes were rinsed four times in either Cook, Vitrolife or KSOM and then cultured in 40-µl drops of the same medium for 4 hours at 37°C and 5% CO_2_. After this short different incubation medium, eggs with two pronuclei (2PNs) were further cultured in KSOM until they were transferred into pseudo-pregnant recipients at one or two-cell stage (Figure 1).

**Figure 1.**
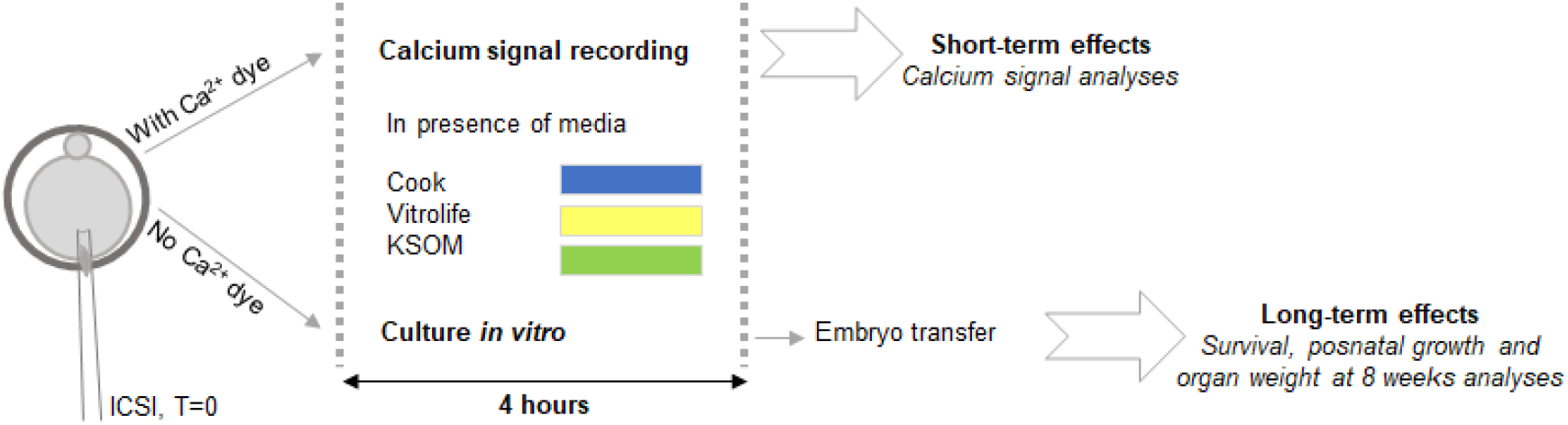
Schematic experimental design. Oocytes were fertilized by ICSI with Ca^2+^ dye for Ca^2+^ recording or without dye for in vitro culture. After ICSI with dye, Ca^2+^ responses were recorded in a microfluidic device until Ca^2+^ oscillations ceased. Eggs were examined for the presence of 2PNs. Ca^2+^ recordings were then analyzed. After ICSI without dye, ICSI-fertilized oocytes are immediately placed in one of the three culture media and cultured for 4 hours at 37°C in the incubator. Eggs with 2PNs were transferred into female recipients at the 1-cell stage as a priority or at 2-cell stage, depending on the availability of recipients. The survival and growth rate of the animals was recorded until 8 weeks of age.

### 2.4. Ca^2+^ signal recording

For Ca^2+^ signal recordings, sperm injections were performed in ICSI medium containing 500 μM Fura-2 dextran (F3029, Thermo Fisher). After injection, two ICSI-fertilized oocytes were placed in a microfluidic chamber mounted on a Nikon TE2000 (NIKON, Japan) inverted microscope equipped with a Fluor 40× oil immersion objective and an EMCCD Photometrics camera (EVOLVE, Roper Scientific, USA). The ICSI-fertilized oocytes were first washed with a rapid flow of KSOM medium at approximately 10 μL s^−1^ to remove residual ICSI medium, followed by stabilization at a slower medium flow rate of 1.4 μL s^−1^ to establish baseline fluorescence measurements in the microfluidic chamber^30^. The KSOM medium was then replaced with one of the three experimental culture media. Intracellular Ca^2+^ signals were monitored using fluorescence excitation provided by a monochromator at alternating wavelengths of 340 nm and 380 nm, with emission recorded at 510 nm. The intracellular Ca^2+^ concentration was expressed as the fluorescence ratio (F340/F380) and recorded at a sampling frequency of 0.5 Hz, beginning 1 min after the ICSI-fertilized oocytes were placed in the microfluidic chamber and continuing until the Ca^2+^ oscillations ceased. Stable experimental conditions were maintained during recording by a constant medium flow, a temperature of 37°C, and regulated gas concentrations. Upon completion of Ca^2+^ recordings, eggs were observed to confirm PN formation.

### 2.5. Embryo transfer

Female mice (RjOrl:SWISS, 8-15 weeks old, Janvier Labs, France) were used as recipients. These females were mated with vasectomized males, and on the morning of the experimental day, they were examined for the presence of a vaginal plug. Eggs were then transferred in groups of 8 into the left oviduct of the recipient females, either at the one-cell stage or the two-cell stage depending on the availability of recipients.

### 2.6. Postnatal phenotype

The postnatal phenotype of the offspring was assessed through weight monitoring, with the animals being weighed weekly starting from the first week after birth. To allow for longitudinal follow-up of each individual, pups were tattooed on their paws during the first postnatal week and weighed weekly in the morning until week eight, whenever on the same weekday as their birth. Offspring from each culture condition remained with their recipient mothers until weaning. All animals were housed in the same room under standardized conditions: A 12:12-hour light/dark cycle was used, and the temperature and humidity were controlled. The animals had ad libitum access to standard chow and water. At the 4th week, the pups were weaned, and males and females were separated, with groups of up to five animals housed in individual cages. Sex ratios were assessed across litters and did not differ significantly between culture conditions (data not shown). At 8 weeks of age, the animals were euthanized by cervical dislocation. For phenotypic analyses, the number of animals dissected at 8 weeks of age was as follows: 15 females and 16 males in the Cook group, 13 females and 14 males in the Vitrolife group, and 13 females and 10 males in the KSOM group. The liver, kidneys, heart, lungs, brain, and fat were then dissected and weighed.

### 2.7. Statistical analysis

Statistical analyses were performed using R software (version 4.3.1; https://www.R-project.org). Figures and graphs were generated using the ggplot2 package (version 3.5.1; https://ggplot2.tidyverse.org). Group differences were evaluated with the Kruskal-Wallis test, a non-parametric method appropriate for comparing multiple independent groups. Pairwise comparisons between groups were conducted using the Dunn test, with Bonferroni correction applied to account for multiple comparisons. Additionally, for Table 3, the chi-square (χ^2^) test was used to analyze contingency tables and compare proportions as specified in the legend. All values are reported as mean ± standard deviation (SD), and detailed statistical tests are indicated in the figure legends and tables. Most of the results derived from KSOM media have been previously published by Ozil et al.^18^.

**Table 3:** Development to term of embryos after ICSI eggs cultured in three different media. The rates of pregnancy, survival to term, and pup survival were not significantly different among the three groups, as determined by Chi-square test. Additionally, the mean litter sizes at birth did not differ significantly between groups, based on Dunn’s post-hoc test with Bonferroni correction.

| Media | No. eggs transferr ed | No. pregnant/N o. recipient | No. pups from eggs transferred in pregnant | No. live pups | Litter size | Ratio F/M |
| --- | --- | --- | --- | --- | --- | --- |
| Cook | 184 | 14/23 (61%) | 35/112(31%) | 32/35(91%) | 2.5±1.4 | 1.00 (16/16) |
| KSOM | 200 | 16/25 (64%) | 42/128(33%) | 41/43(95%) | 2.7±2 | 0.86 (18/21) |
| Vitrolif e | 152 | 10/19 (53%) | 32/80(40%) | 27/32(84%) | 3.2±1 | 0.93 (13/14) |

## 3. RESULTS

### 3.1. Impact of culture media composition on Ca^2+^ responses in mouse ICSI-fertilized oocytes

The composition of three media used for in vitro embryo culture was compared (Table 1). The media differ primarily in the ratio of the two main energy sources for early embryos, lactate and pyruvate, as well as in their amino acid content. Specifically, Cook contains both essential and non-essential amino acids, while Vitrolife contains only essential amino acids, and KSOM lacks amino acids entirely. In addition, potassium and magnesium concentrations are higher in Cook and Vitrolife than in KSOM.

To assess the impact of these different media on Ca^2+^ responses in oocytes fertilized by ICSI, we evaluated several key parameters. These parameters included the Ca^2+^ oscillation patterns observed in vitro, the total number of calcium peaks over four hours, the duration of each Ca^2+^ oscillation event, and its frequency over one hour.

Typical individual Ca^2+^ responses following oocyte fertilization in Cook, Vitrolife or KSOM media are shown in Figure 2. Time zero corresponds to the start of calcium recording in the microfluidic chamber, initiated 2-3 minutes after ICSI. In each response, the first oscillation was characterized by a series of super-oscillations and lasted longer than subsequent ones, as described by Ozil *et al.*^17^. This initial oscillation was followed by repeated oscillations that could be recorded for several hours. In particular, the number of Ca^2+^ oscillations (Ca^2+^ peaks) was lower in the Cook and Vitrolife media compared to the KSOM medium (Figure 2).

**Figure 2.**
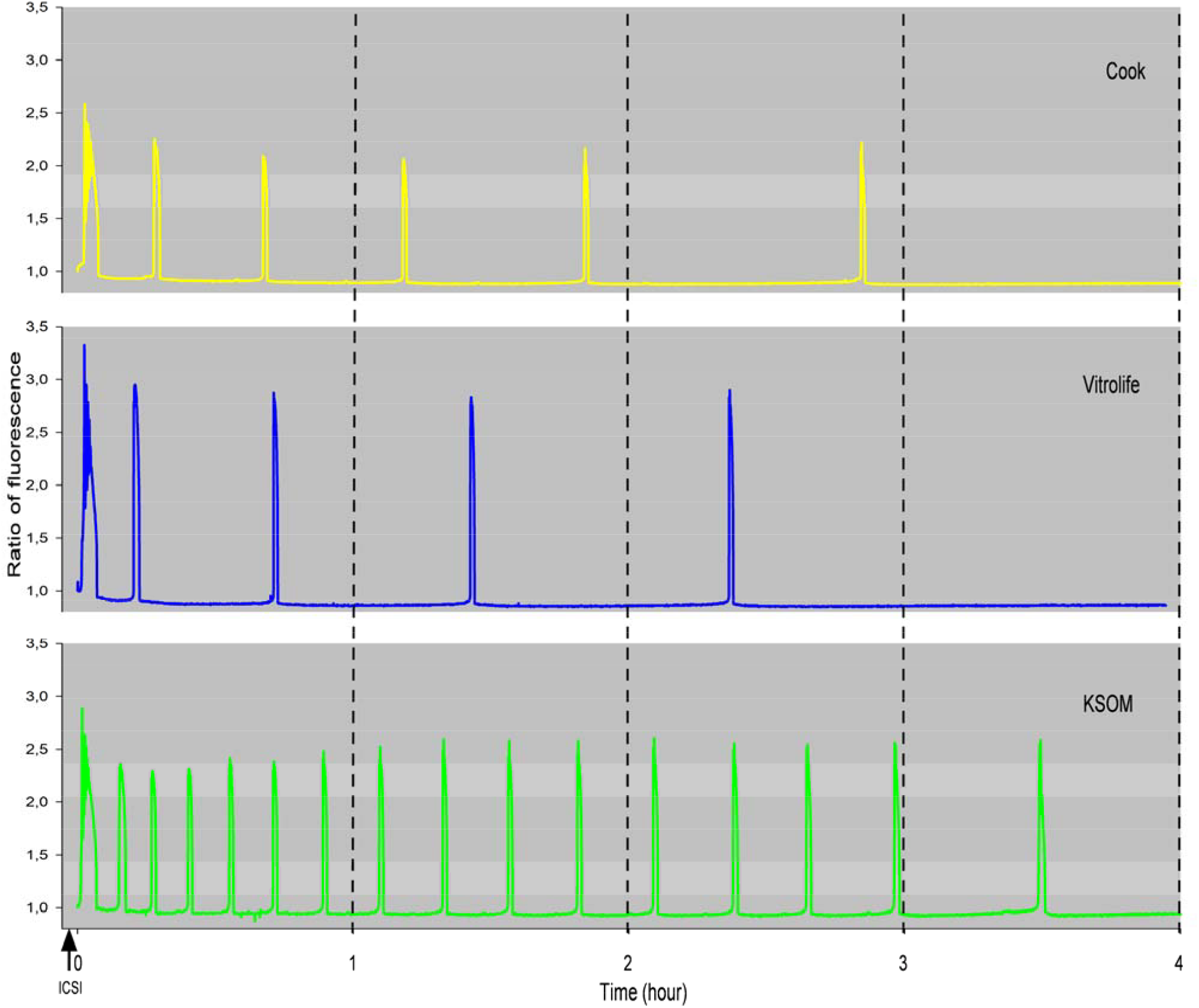
Representative individual Ca^2+^ responses in ICSI-fertilized oocytes cultured in three media for 4 hours. Typical Ca^2+^ responses are shown in Cook (yellow with 6 Ca^2+^oscillations), Vitrolife (blue with 5 Ca^2+^oscillations), and KSOM (green with 16 Ca^2+^oscillations) media. Time zero corresponds to the start of Ca^2+^ recording in the microfluidic chamber, initiated within 2-3 minutes following ICSI.

Compilation of multiple recordings revealed that the mean number of Ca^2+^ oscillations in ICSI-fertilized oocytes was three times lower in Cook or Vitrolife than in KSOM, 5.6 ± 1.6 and 5.1 ± 1.6 vs. 15.6 ± 5.9, respectively (P < 0.05; Figure 3A). There was no difference between Cook and Vitrolife. The dispersion of the number of oscillations is lower in Cook or Vitrolife compared to the KSOM medium (Figure 3A). Furthermore, the mean number of Ca^2+^ oscillations within the first hour was twice lower in Cook or Vitrolife than in KSOM medium, 3.1 ± 0.8 vs. 6.4 ± 2.4, respectively (P < 0.05; Figure 3B). This significant difference was even more pronounced over the following hours: threefold in the second and third hours, and sevenfold in the fourth hour (P < 0.05; Figure 3C through 3E).

**Figure 3.**
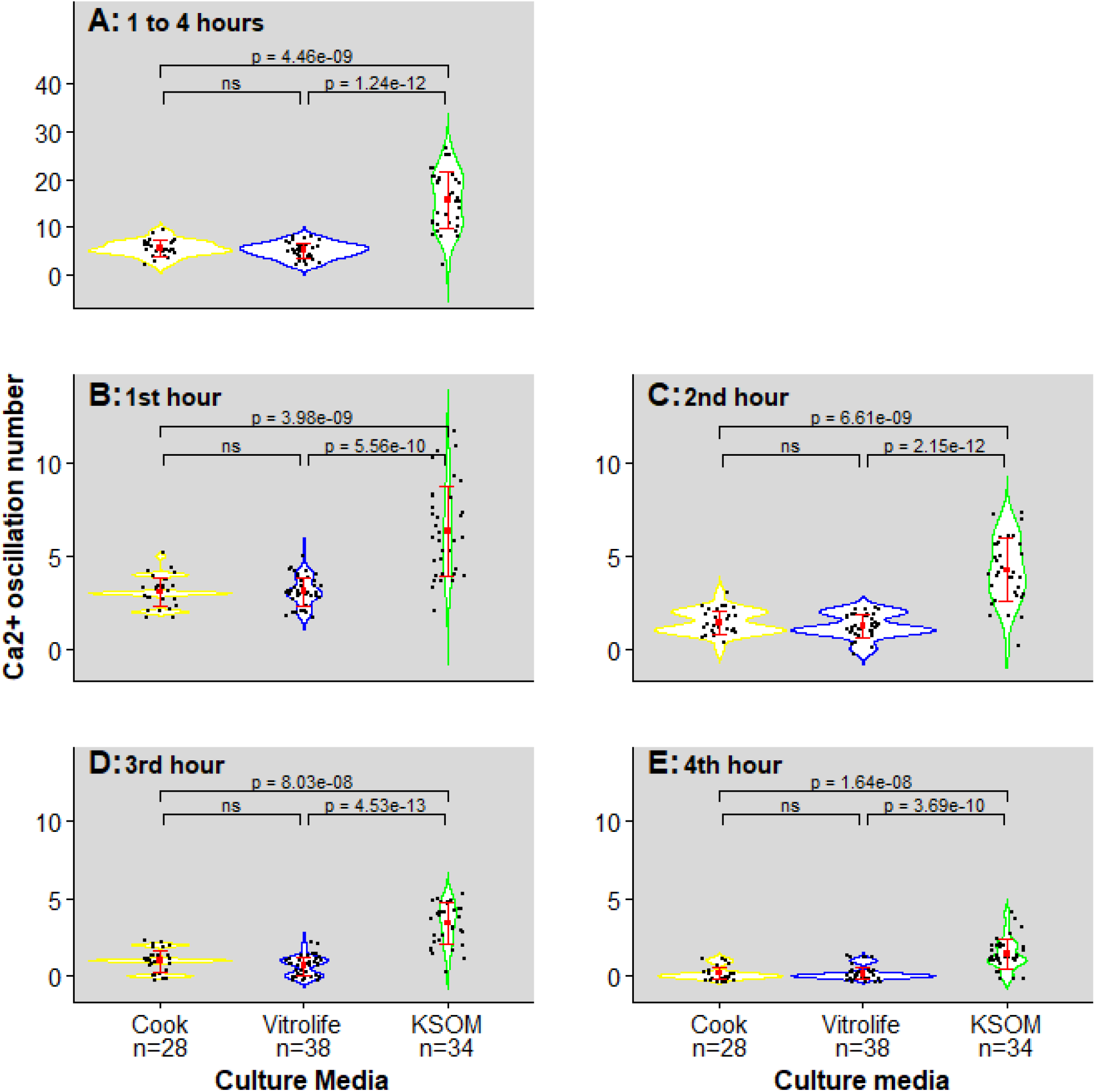
Number of Ca^2+^ oscillations in ICSI-fertilized oocytes cultured in three media. Figure 3A shows the mean number of Ca^2+^ responses for ICSI-fertilized oocytes cultured for 4 hours in Cook, Vitrolife or KSOM media. Each black dot represents a complete Ca^2+^ record, from the initial Ca^2+^ peak to the cessation of oscillations. Figures 3B through 3E show the mean number of Ca^2+^ responses from hour 1 through hour 4 for ICSI-fertilized oocytes cultured in Cook, Vitrolife, or KSOM media. The mean number of responses for each medium is indicated by a red square, and the standard deviation is indicated by the error bars in red. P-values obtained by Dunn’s post-hoc test (with Bonferroni correction) are shown. Statistical significance is indicated by horizontal brackets and p-values, with significant differences (p < 0.05) indicated and non significant comparisons marked “ns”.

In addition to the lower number of oscillations in Cook and Vitrolife compared to the other medium, the total duration of the ICSI-fertilized oocyte response was also reduced, as was the frequency of oscillations (Figure 4). The mean durations in Cook and Vitrolife were 2 hours, 26 minutes, and 59 seconds and 2 hours, 5 minutes, and 5 seconds, respectively, compared to 3 hours, 17 minutes, and 46 seconds for KSOM. The mean frequencies of Ca^2+^ responses in Cook and Vitrolife were 4.7 ± 1.2 and 2.5 ± 1.0, respectively, vs. 2.7 ± 1.0 in KSOM. However, there was no difference between Cook and Vitrolife in either duration or frequency.

**Figure 4.**
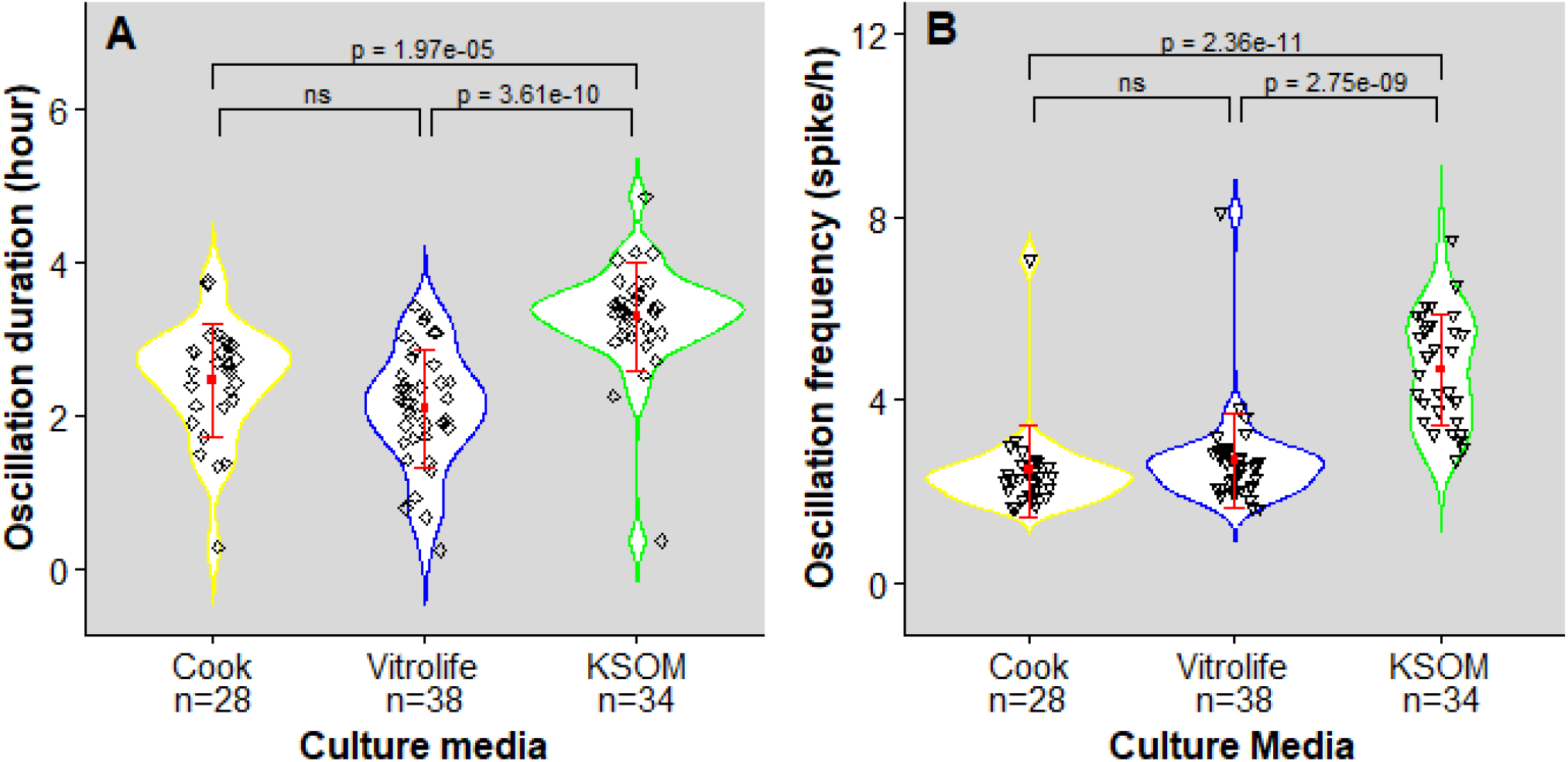
Duration and frequency of Ca^2+^ oscillations in ICSI-fertilized oocytes cultured in three media. 4A shows the duration and 4B the frequency of Ca^2+^ responses for ICSI-fertilized oocytes cultured in Cook, Vitrolife, or KSOM media. Each black dot represents a complete Ca^2+^ record, from the initial Ca^2+^ peak to the cessation of oscillations. The mean duration of responses for each medium is indicated by a red square, with the standard deviation represented by the error bars in red. P-values obtained by Dunn’s post-hoc test (with Bonferroni correction) are shown. Statistical significance is indicated by horizontal brackets and p-values, with significant differences (p < 0.05) indicated and non significant comparisons marked “ns”.

### 3.2. Development to term and postnatal growth

After ICSI, embryos were cultured in the three different media for four hours until 2PNs were formed. They were then transferred to recipient females (Table 3). The rate of term development was 31 to 40%, with no statistical difference between the three media. A few pups died during the first week. Thereafter, the growth of male and female pups was followed through week 8 by measuring their weight once a week.

The rates of pregnancy (53 to 64 %), survival to term (31 to 40%) and pup survival (84 to 95%) were not statistically different among the three groups according to the chi-squared test. In addition, mean litter size at birth was not significantly different between groups Cook (2.5 ± 1.4), KSOM (2.7 ± 2), and Vitrolife (3.2 ± 1) as assessed by Dunn’s post-hoc test with Bonferroni correction (Table 3). Regarding the sex ratio (0.86 to 1), no significant differences were observed in the proportions of females and males across the groups, as determined by the chi-squared test (Table 3).

### 3.3. Postnatal growth

#### 3.3.1. Growth differences in pups from ICSI eggs cultured in KSOM compared to Cook and Vitrolife media

Pups were weighed one week after birth and then for eight weeks as the mice grew to adulthood. No differences were observed between the mean growth curves of ‘Cook’ and ‘Vitrolife’ females and males. The mean growth curve of ‘Cook’ females was significantly higher than that of ‘KSOM’ females at every time point (P < 0.001; Figure 5A), although there were two individual growth curves that were very low in the Cook medium (Figure S1A). The mean growth curve of ‘Vitrolife’ females was significantly higher than that of ‘KSOM’ females from week 2 to week 8 (P < 0.05; Figure 5A). The same significant difference is observed for males, but with a less pronounced effect, with ‘KSOM’ males having a lower weight than those in the other two media at most time points, except for week 1 in Cook and weeks 3-5 in Vitrolife (P < 0.05; Figure 5B). There was an individual growth curve for the male that was very low in the Cook medium (Figure S1B).

**Figure 5.**
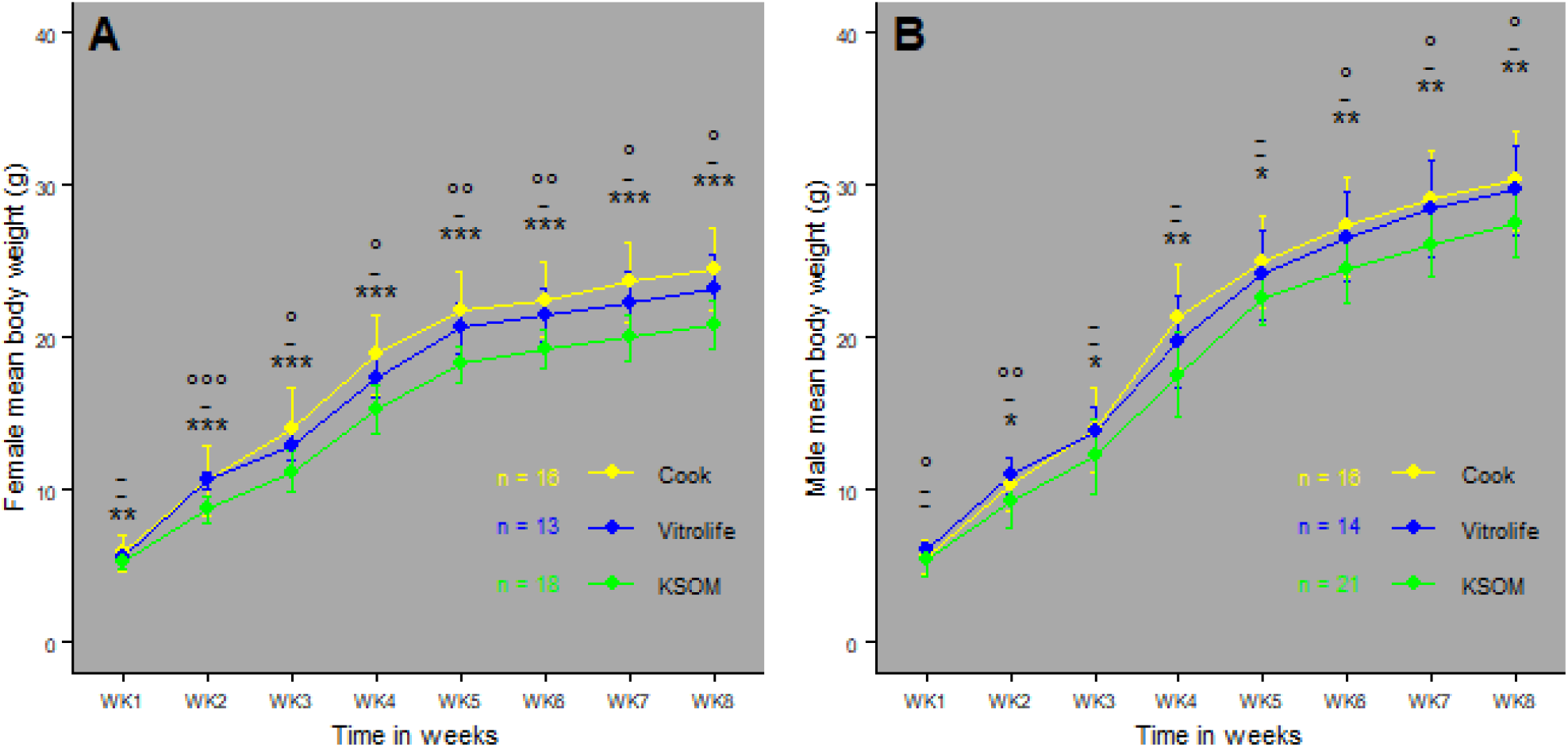
Postnatal growth of pups up to 8 weeks of age. 5A and 5B show the mean growth curve of females (A) and males (B). Cook’s growth profile is shown in the yellow line, Vitrolife in the blue line, and KSOM in the green line. Error bars represent standard deviations. Pairwise comparison of animal weight in different media from 1 to 8 weeks of age. Statistical analyses were performed separately for females and males using Dunn’s post-hoc test, with Bonferroni correction applied to all pairwise comparisons for each week. The p-values resulting from these comparisons are indicated by different labels. Comparison KSOM - Vitrolife: ° p ≤ 0.05, °° p ≤ 0.01, °°° p ≤ 0.001. Comparison Cook - Vitrolife : ¤ p ≤ 0.05, ¤¤ p ≤ 0.01, ¤¤¤ p ≤ 0.001.Comparison Cook - KSOM : * p ≤ 0.05, ** p ≤ 0.01, *** p ≤ 0.001.

### 3.4. Organ and tissue weights of mice at the 8th week of age

#### 3.4.1. Differences in specific organ-to-body weight ratios in female offspring

At 8 weeks of age, Cook and Vitrolife female mice had significantly higher organ weights, including brain, heart, kidney, liver, and lung, compared to KSOM female mice, as shown in Figure S2. They also had greater fat deposition around the kidney and gonads. After normalization for body weight, Cook and Vitrolife females had a higher kidney/body weight ratio than KSOM females. Cook females had a higher white fat to body weight ratio and a lower brain to body weight ratio than KSOM females. Vitrolife females had a higher liver-to-body weight ratio than KSOM females, as shown in Figure 6.

**Figure 6.**
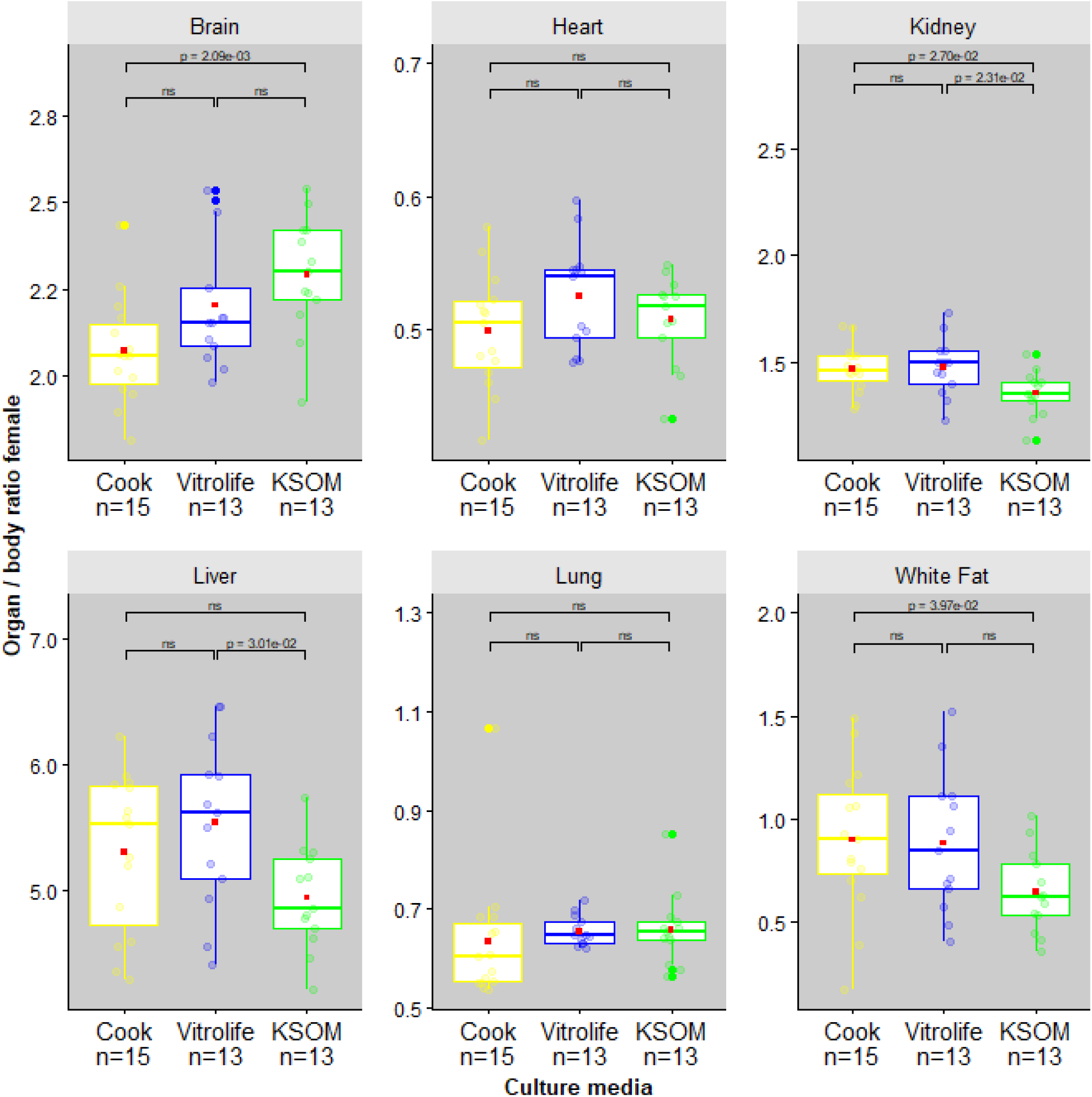
Mean organ/body weight ratios of female offspring at 8 weeks of age. Box plots illustrate the distribution of organ and tissue weight ratios for each group. The red square in each box represents the mean ratio, while jittered round colored dots indicate individual data points. Statistical significance is indicated by horizontal brackets and p-values, with significant differences (p < 0.05) indicated and non significant comparisons marked “ns”.

#### 3.4.2. No differences in specific organ-to-Body weight ratios among male offspring

At 8 weeks of age, Cook and Vitrolife male mice had significantly higher absolute organ weights for brain and heart compared to KSOM male mice, as shown in Figure S3. However, only Cook male mice had a significantly higher absolute kidney weight compared to KSOM. After normalization for body weight, no significant differences in organ-to-body weight ratios were observed between the male groups, as shown in Figure 7.

**Figure 7.**
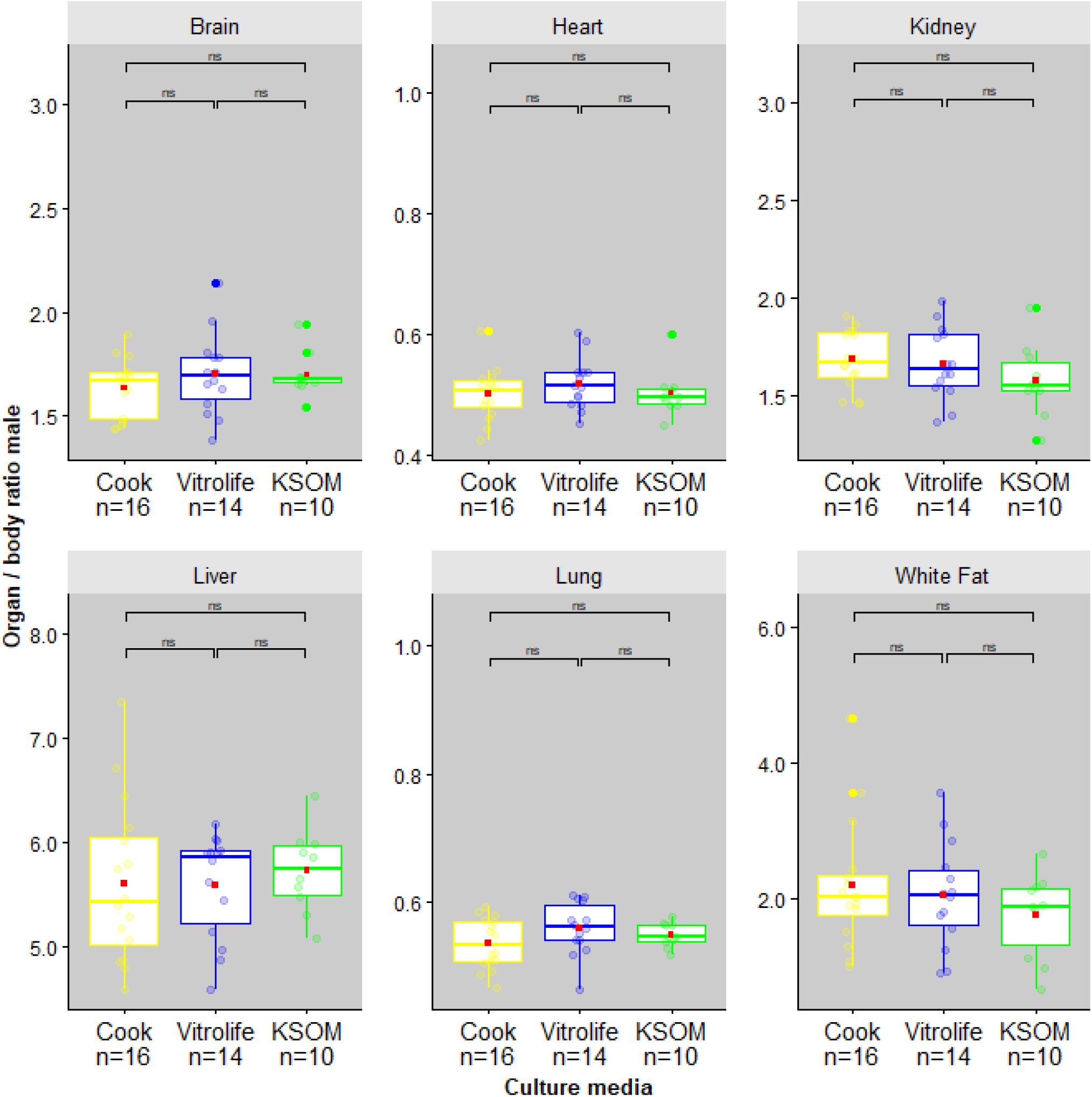
Mean organ-to-body weight ratios of male offspring at 8 weeks of age. Box plots illustrate the distribution of organ and tissue weight ratios for each group. The red square in each box represents the mean ratio, while jittered round colored dots indicate individual data points. Statistical significance is indicated by horizontal brackets and p-values, with significant differences (p < 0.05) indicated and non significant comparisons marked “ns”.

## 4. DISCUSSION

Sperm entry during fertilization triggers a rapid increase in intracellular free Ca^2+^ concentration in the oocyte. This Ca^2+^ spike initiates a cascade of physiological and biochemical changes that activate the developmental program of the early embryo^5^. Our previous studies demonstrated that the composition of two mouse embryo culture media, M16 and KSOM—both containing the same chemical components in different proportions—modulates the pattern of calcium oscillations, ultimately influencing postnatal growth^18^.

Despite the increasing number of children born worldwide through ICSI, the impact of ART culture media composition on the metabolic activity of ICSI-fertilized oocytes and its long-term effects after embryo transfer remain largely unexplored in mice. Investigating these interactions at the earliest developmental stage immediately after fertilization and over a brief 4-hour period, corresponding to Ca^2+^ oscillations and PN formation could provide new insights into how culture media influence embryonic metabolism. These findings could ultimately contribute to optimizing assisted reproductive technology protocols.

Determining whether culture conditions at the time of fertilization influence phenotypic characteristics, such as body weight, is crucial. If proven, this would support the hypothesis developed by Ducibella^31^ that the embryonic environment from fertilization onward induces metabolic changes with long-term consequences, aligning with the Developmental Origins of Health and Disease (DOHaD) theory^32^.

In this study, we compared three culture media, including two commonly used in ART, and identified distinct Ca^2+^ response patterns immediately after fertilization. These differences correlated with variations in postnatal growth and organ weight. Specifically, our findings reveal that the two ART media induce a different short-term response and long-term developmental outcomes compared to KSOM.

### Ca^2+^ response

Our results confirm and extend previous findings in mice^18^, showing that sperm-induced Ca^2+^ oscillations are more efficiently generated in the presence of a low Mg^2+^/high Ca^2+^ culture medium, such as KSOM. Collectively, these data indicate that the external [Mg^2+^]/[Ca^2+^] ratio plays a key role in shaping the pattern of Ca^2+^ oscillations by regulating Ca^2+^ influx. Modifying this ratio using different culture media—KSOM, Cook, and Vitrolife—during egg activation significantly impacts the Ca^2+^ response. KSOM, containing 0.2 mM Mg^2+^ and 1.7 mM Ca^2+^, promotes a distinct oscillatory pattern compared to Cook and Vitrolife, which contain 1.5 mM and 1.7 mM Mg^2+^ with 1.1 mM Ca^2+^, respectively^22^. These results confirm that Mg^2+^ influence Ca^2+^ influx, thereby modulating oscillation frequency and amplitude^18^.

During ICSI, sperm-specific phospholipase C zeta (PLCζ) hydrolyzes phosphatidylinositol 4,5-bisphosphate, generating inositol trisphosphate (IP3) and diacylglycerol^33^. IP3 then binds to its receptors on the endoplasmic reticulum, triggering the release of stored Ca^2+^ into the cytosol, initiating a series of oscillations lasting several hours^34^. However, sustaining these oscillations requires Ca^2+^ influx across the oocyte plasma membrane^35^, primarily mediated by CaV3.2 and TRPM7 channels^36^. Notably, TRPM7 also plays a role in Mg^2+^ homeostasis, potentially competing with Ca^2+^ influx^37^.

Our study confirms that higher Mg^2+^ concentrations reduce both the frequency and variability of Ca^2+^ oscillations, whereas lower Mg^2+^ levels permit a broader oscillatory range with greater interindividual variability as previously shown^18^. This pattern is evident when comparing Cook/Vitrolife with KSOM.

Another key difference between KSOM and the ART media (Cook/Vitrolife) lies in the type and concentration of added proteins—BSA in KSOM versus HSA in Cook and Vitrolife, with the latter containing a markedly higher concentration (5 mg/mL vs. 1 mg/mL). However, experiments using Global medium—which also contains 5 mg/mL of HSA but has a low [Mg^2+^]/[Ca^2+^] ratio (0.2/1.9 mM)—confirmed that differences in Ca^2+^ oscillatory patterns persisted (unpublished data). These findings confirm that the Mg^2+^/Ca^2+^ and [Ca^2+^]? summation^28^ play a crucial role as determinants of oscillatory dynamics.

Furthermore, variations in lactate and pyruvate concentrations, as well as the lactate/pyruvate ratio, differentiate KSOM from Cook/Vitrolife. However, despite Global medium having a lower lactate/pyruvate ratio^21^, it still induces a higher Ca^2+^ response than Vitrolife, reinforcing the conclusion that ionic composition —particularly the Mg^2+^/Ca^2+^ ratio— is an important factor in the regulation of Ca^2+^ oscillations at this stage.

Thus, modulating extracellular Mg^2+^ concentration directly influences the dynamics of Ca^2+^ oscillations post-fertilization. Careful control of this balance in embryo culture media is therefore crucial for optimizing ART protocols^18^.

### Growth Rates and Organ/Tissue Weights

Since Ca^2+^ oscillations play a key role in initiating embryonic development, alterations in their pattern could have long-term consequences. To explore this, we analyzed postnatal growth and organ weights in mice cultured in different media.

Our results reveal significant effects of the culture media on growth in both males and females, aligning with previous mouse studies and human IVF reports. Even a brief exposure to specific media between fertilization and PN formation can influence postnatal development, from birth to adulthood.

Notably, females from the Cook and Vitrolife groups exhibited increased organ and tissue weights, while in males, differences were restricted to three organs: the brain, heart, and kidney. This suggests a sex-specific impact of culture media on organ development. After normalization by body weight, females from Cook and Vitrolife showed an increased kidney and fat ratio, while the brain ratio decreased. While these analyses revealed some sex- and organ-specific tendencies, no consistent or significant effect of culture medium on disproportionate organ growth was identified. Limited sample size and statistical power constrain mechanistic interpretation. Further studies will be needed to determine whether the observed trends reflect targeted developmental programming or broader somatic growth effects.

Strikingly, we observed significant differences between Cook, KSOM, and Vitrolife after only 4 hours of culture. These differences persisted into adulthood, affecting both body weight and organ/body weight ratios in female mice. This underscores the potential for even a short exposure to specific culture conditions to induce long-lasting developmental effects.

Similar trends have been reported in humans: embryos cultured in Vitrolife medium for 2–3 days had significantly higher birth weights than those cultured in Cook medium^8^. Follow-up studies confirmed that these differences persisted into early childhood, with children from the Vitrolife group exhibiting higher weight, height, BMI, waist circumference, and truncal adiposity at age nine^10,11^.

A mechanistic metabolic link has been recently discovered between abnormal calcium exposure at fertilization and the long-term health of offspring, which has significant implications for ART. In a 2025 study, Savy et al. demonstrated that excessive Ca^2+^ signals at fertilization disrupt mitochondrial metabolism in mouse embryo^38^. This disruption increases pyruvate dehydrogenase activity and acetyl-CoA production while reducing lactate level. These metabolic shifts alter the availability of key epigenetic substrates, notably affecting histone modifications such as H3K27 acetylation and H3K18 lactylation during the transition from the 1-cell to 2-cell stage. This epigenetic reprogramming impairs embryonic genome activation, resulting in long-lasting changes in gene expression, particularly in adipose tissue. These changes contribute to metabolic alterations, such as elevated fasting glucose levels, in adult male offspring. These findings emphasize the critical role of Ca^2+^-driven metabolic signaling in shaping the epigenetic landscape at the beginning of development and the importance of carefully considering calcium manipulation strategies in ART protocols^31^.

Taken together, our findings reinforce the profound and lasting impact of embryo culture conditions on postnatal growth and metabolic development. Understanding these mechanisms is essential to refine ART protocols and improve long-term health outcomes for individuals conceived through assisted reproduction. In line with this perspective, a seminal study focused on the systemic modulation of calcium dynamics, egg metabolism and long-term consequences through IVF media formulation (J.-P. Ozil, et al., in preparation).

## 5. CONCLUSION

This study confirms previous studies that a 4-hour post-fertilization exposure of ICSI-fertilized oocytes to different culture media influences both early Ca^2+^ signaling and long-term development in mice. Cook and Vitrolife media, with high [Mg^2+^]/[Ca^2+^] ratios, generate distinctCa^2+^activation. These early differences result in significant phenotypic variations in adulthood,highlighting the critical influence of the short window between ICSI and PN formation. However,species-specific differences in embryonic development and epigenetic regulation limit direct extrapolation to humans. Future research should further investigate the interactions between ionic balance, metabolism, and epigenetic modifications to optimize ART culture conditions and improve long-term outcomes.

## AUTHOR CONTRIBUTIONS

B.B. conducted research; B.B. and T.SB. performed research; B.B., T.SB and A.F analyzed data; and B.B. and A.J. wrote the paper.

## ACKNOWLEDGMENTS

The authors would like to thank Dr. Jean-Pierre Ozil for his decisive role in conceiving, designing and writing the initial project, which was funded by the agence de la biomedecine. The subsequent submission and funding of the project was made possible thanks to Bernadette Banrezes. We gratefully acknowledge Dr. J-P Ozil’s commitment and contributions to the scientific rationale, the development of the hypotheses and the design of the experimental framework. We thank Sophie Calderari for critical reading of the manuscript. Finally, we thank the unit of Infectiology of Fishes and Rodents Facility (IERP-UE907, Jouy-en-Josas Research Center: https://doi.org/10.15454/1.5572427140471238E12; member of the National Infrastructure EMERG’IN), for excellent animal care and help with the animal experiments.

## ETHICS STATEMENT

The animal experiments were approved by the French Ministry of Higher Education, Research and Innovation and by the local Animal Experiments Ethics Committee of the INRAE (approval nos. 12-165 and 16-22). All animal procedures were performed in accordance with the Code for Methods and Welfare Considerations in Behavioral Research (Directive 2010/63/EU).

## Supporting Information

**Figure S1.**
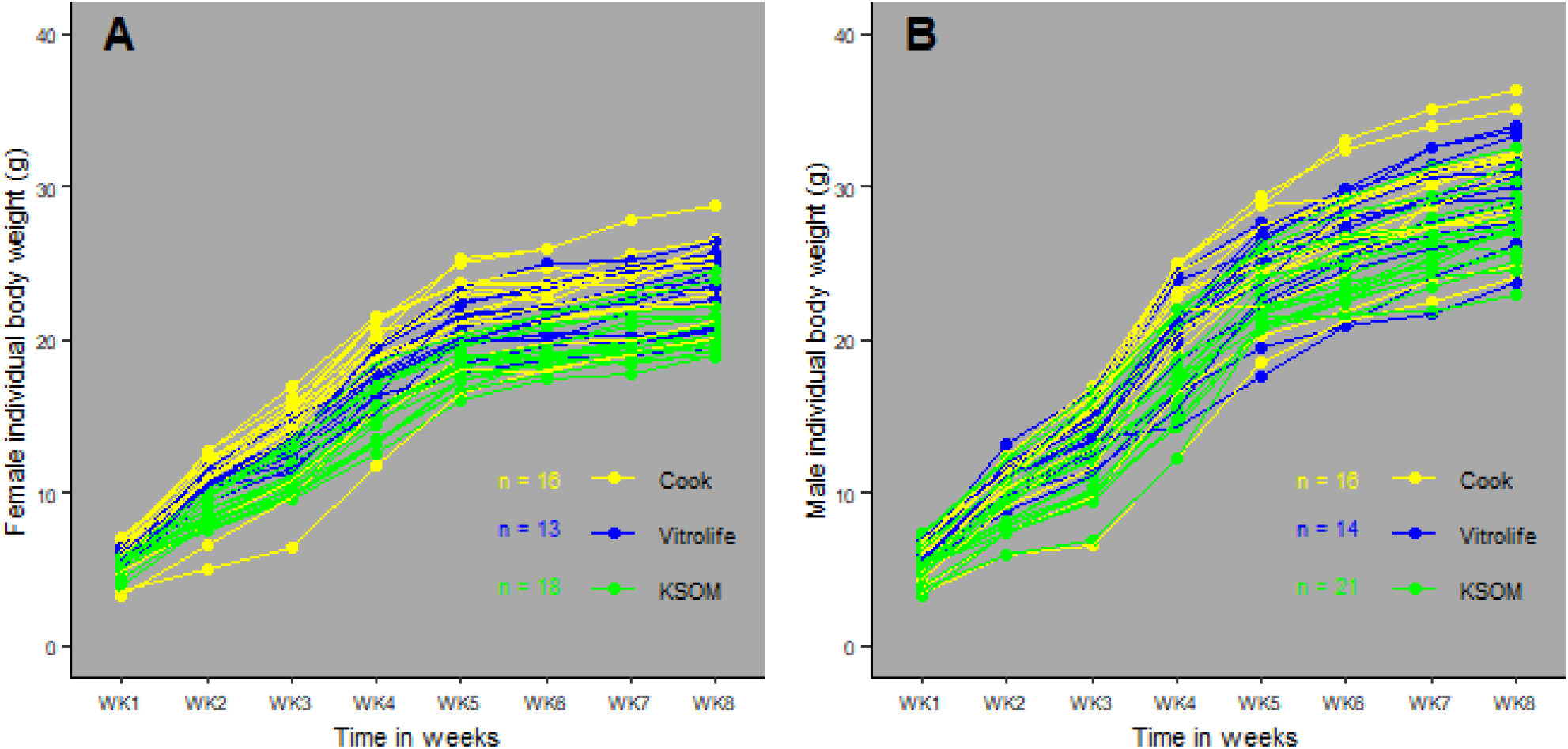
Postnatal growth of pups up to 8th week of age. 1A and 1B show individual growth curve of females (A) and males (B). Cook’s growth profile is shown in the yellow line, Vitrolife’s in the blue line and KSOM’s in the green line.

**Figure S2.**
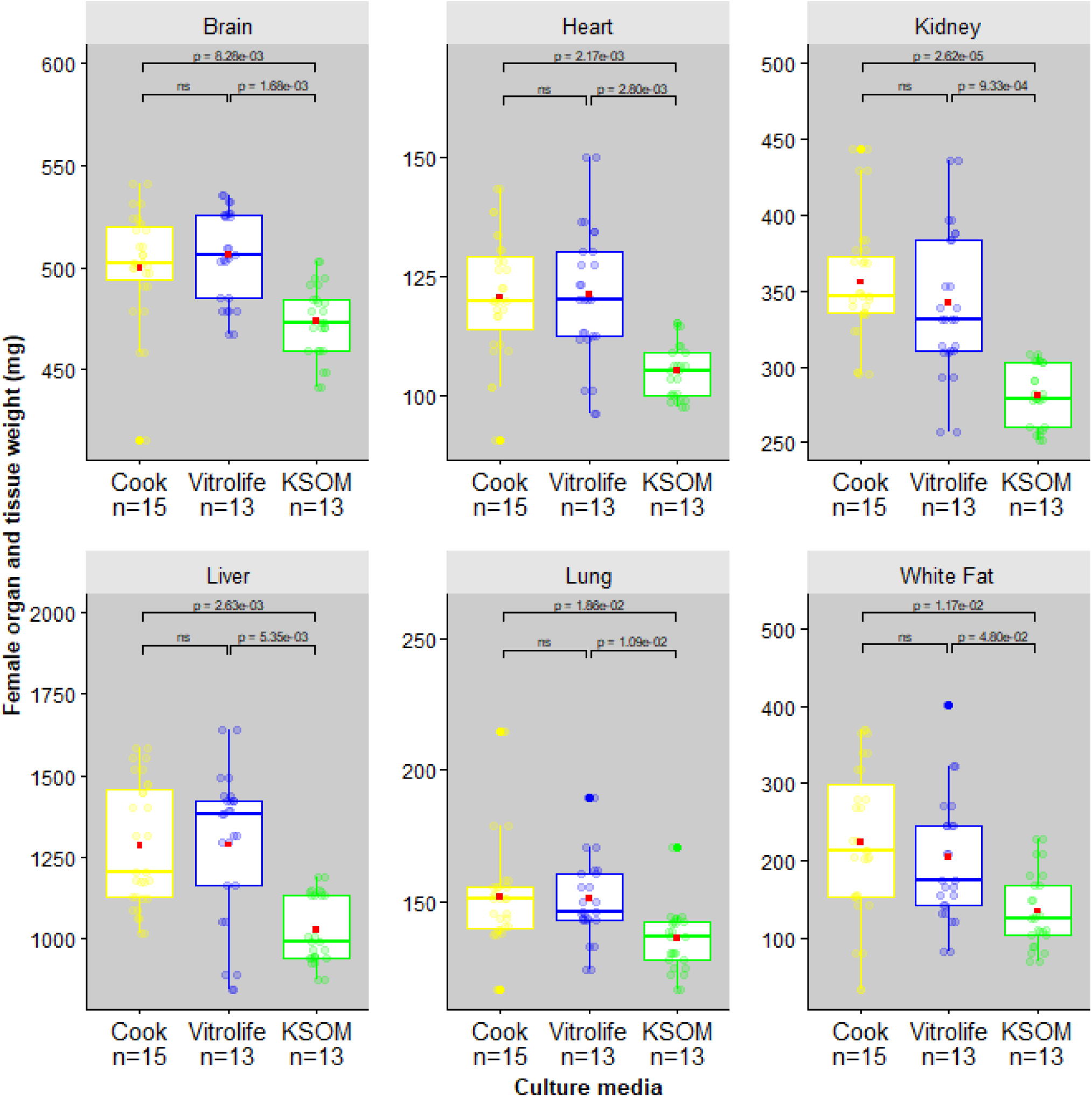
Mean organ and tissue weights of female offspring at 8 weeks of age. Box plots illustrate the distribution of organ and tissue weights for each group. The red square in each box represents the mean organ weight, while jittered round colored dots indicate individual data points. Statistical significance is indicated by horizontal brackets and p-values, with significant differences (p < 0.05) marked and nonsignificant comparisons marked “ns”. The figure shows that Cook and Vitrolife females had significantly higher organ weights (brain, heart, kidney, liver, and lung) and greater fat deposition (white fat around the kidney and gonads) compared to KSOM females.

**Figure S3.**
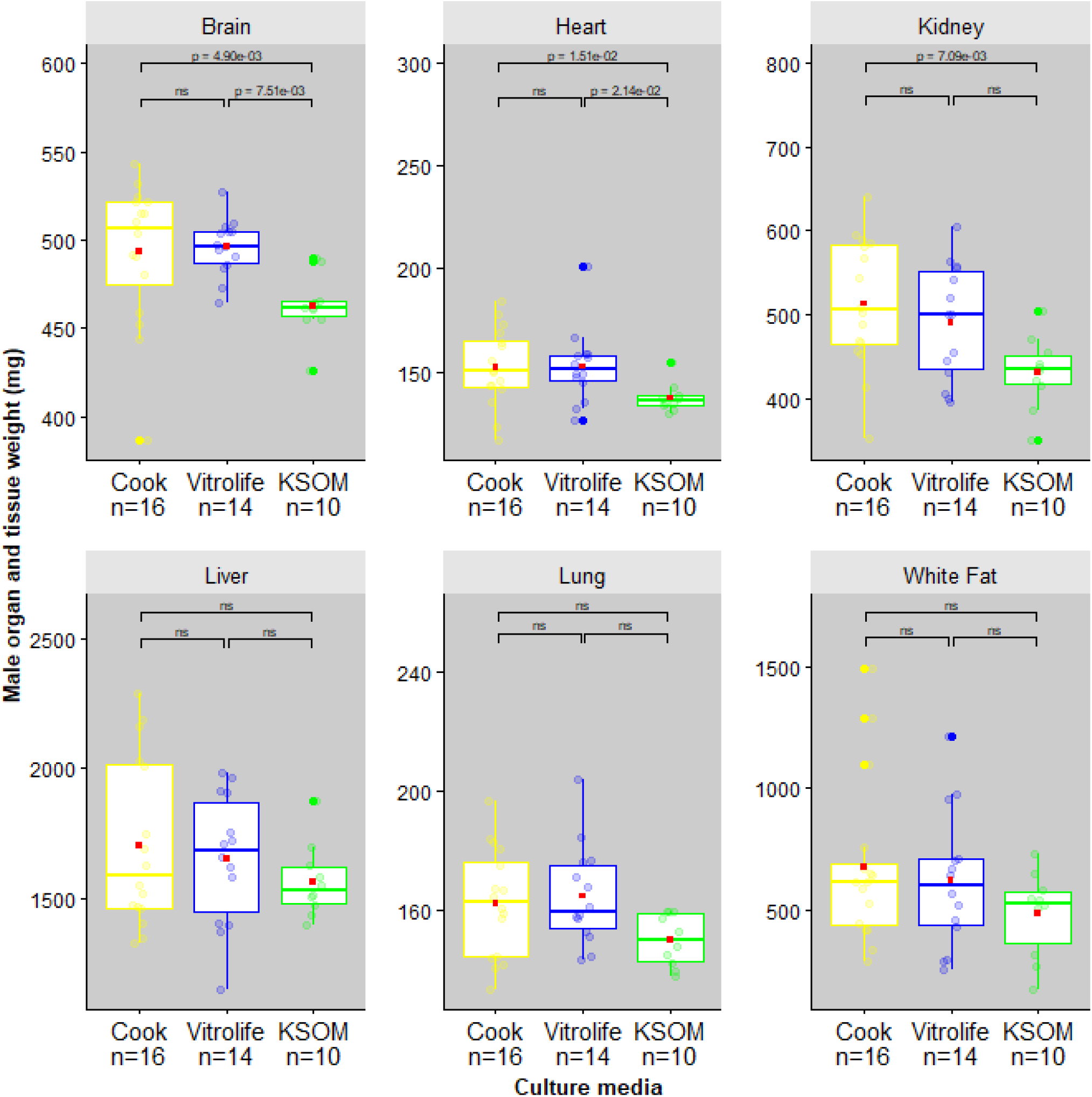
Mean organ and tissue weights of male offspring at 8 weeks of age. Statistical significance is indicated by horizontal brackets and p-values, with significant differences (p < 0.05) marked and nonsignificant comparisons marked “ns”. The figure shows that while Cook and Vitrolife males had significantly higher brain and heart weights than KSOM males, only Cook males had a significantly higher kidney weight. Additionally, both Cook and Vitrolife males exhibited greater fat deposition (white fat around the kidney and gonads). Statistical significance is indicated by horizontal brackets and p-values, with significant differences (p < 0.05) indicated and non significant comparisons marked “ns”.

## REFERENCES

1. Smeenk J, Wyns C, De Geyter C, Kupka M, Bergh C, Cuevas Saiz I, et al. ART in Europe, 2019:results generated from European registries by ESHRE. Hum Reprod. 2023;38(12):2321–38.

2. Sunderam S, Zhang Y, Jewett A, Kissin DM. State-Specific Assisted Reproductive Technology Surveillance, United States: 2019 data brief. Atlanta (GA): Centers for Disease Control and Prevention; 2021 Oct 10.

3. Sunderam S, Kissin DM, Zhang Y, Jewett A, Boulet SL, Warner L, et al. Assisted Reproductive Technology Surveillance United States, 2018. MMWR Surveill Summ. 2022;71(4):1–19.

4. Dumollard R, Marangos P, Fitzharris G, Swann K, Duchen M, Carroll J. Sperm-triggered [Ca2+] oscillations and Ca2+ homeostasis in the mouse egg have an absolute requirement for mitochondrial ATP production. Development. 2004;131(13):3057–67.

5. Machaty Z. The signal that stimulates mammalian embryo development. Front Cell Dev Biol. 2024;12:1474009.

6. Zandstra H, Van Montfoort APA, Dumoulin JCM. Does the type of culture medium used influence birthweight of children born after IVF? Hum Reprod. 2015;30(3):530–42.

7. Sciorio R, Rinaudo P. Culture conditions in the IVF laboratory: state of the ART and possible new directions. J Assist Reprod Genet. 2023;40(11):2591–607.

8. Dumoulin JC, Land JA, Van Montfoort AP, Nelissen EC, Coonen E, Derhaag JG, et al. Effect of in vitro culture of human embryos on birthweight of newborns. Hum Reprod. 2010;25(3):605–12.

9. Nelissen EC, Van Montfoort AP, Coonen E, Derhaag JG, Geraedts JP, Smits LJ, et al. Further evidence that culture media affect perinatal outcome: findings after transfer of fresh and cryopreserved embryos. Hum Reprod. 2012;27(7):1966–76.

10. Kleijkers SHM, Van Montfoort APA, Smits LJM, Viechtbauer W, Roseboom TJ, Nelissen ECM, et al. IVF culture medium affects postnatal weight in humans during the first 2 years of life. Hum Reprod. 2014;29(4):661–9.

11. Zandstra H, Brentjens LBPM, Spauwen B, Touwslager RNH, Bons JAP, Mulder AL, et al. Association of culture medium with growth, weight and cardiovascular development of IVF children at the age of 9 years. Hum Reprod. 2018;33(9):1645–56.

12. Carrasco B, Boada M, Rodríguez I, Coroleu B, Barri PN, Veiga A. Does culture medium influence offspring birth weight? Fertil Steril. 2013;100(5):1283–8.

13. Wunder D, Ballabeni P, Roth-Kleiner M, Primi M, Senn A, Chanson A, et al. Effect of embryo culture media on birthweight and length in singleton term infants after IVF-ICSI. Swiss Med Wkly. 2014;140:w14038.

14. Petersen CG, Mauri AL, Vagnini LD, Renzi A, Petersen B, Matilla MC, et al. Randomized comparison of two commercial culture media (Cook and Vitrolife) for embryo culture after IMSI. JBRA Assist Reprod. 2019;23(2):138–43.

15. Vrooman LA, Rhon-Calderon EA, Suri KV, Dahiya AK, Lan Y, Schultz RM, et al. Placental abnormalities are associated with specific windows of embryo culture in a mouse model. Front Cell Dev Biol. 2022;10:884088.

16. Ecker DJ, Stein P, Xu Z, Williams CJ, Kopf GS, Bilker WB, et al. Long-term effects of culture of preimplantation mouse embryos on behavior. Proc Natl Acad Sci U S A. 2004;101(6):1595–600.

17. Banrezes B, Sainte-Beuve T, Canon E, Schultz RM, Cancela J, Ozil JP. Adult body weight is programmed by a redox-regulated and energy-dependent process during the pronuclear stage in mouse. PLoS One. 2011;6(12):e29388.

18. Ozil JP, Sainte-Beuve T, Banrezes B. [Mg2+]o/[Ca2+]o determines Ca2+ response at fertilization: tuning of adult phenotype? Reproduction. 2017;154(5):675–93.

19. Fleming TP, Watkins AJ, Velazquez MA, Mathers JC, Prentice AM, Stephenson J, et al. Origins of lifetime health around the time of conception: causes and consequences. Lancet. 2018;391(10132):1842–52.

20. Ermisch AF, Herrick JR, Pasquariello R, Dyer MC, Lyons SM, Broeckling CD, et al. A novel culture medium with reduced nutrient concentrations supports the development and viability of mouse embryos. Sci Rep. 2020;10(1):9263.

21. Velazquez MA, Idriss A, Chavatte-Palmer P, Fleming TP. The mammalian preimplantation embryo: Its role in the environmental programming of postnatal health and performance. Anim Reprod Sci. 2023;256:107321.

22. Morbeck DE, Krisher RL, Herrick JR, Baumann NA, Matern D, Moyer T. Composition of commercial media used for human embryo culture. Fertil Steril. 2014;102(3):759–66.

23. Morbeck DE, Baumann NA, Oglesbee D. Composition of single-step media used for human embryo culture. Fertil Steril. 2017;107(4):1055–60.

24. Zagers MS, Laverde M, Goddijn M, De Groot JJ, Schrauwen FAP, Vaz FM, et al. The composition of commercially available human embryo culture media. Hum Reprod. 2025;40(1):30–40.

25. Delaroche L, Oger P, Genauzeau E, Meicler P, Lamazou F, Dupont C, et al. Embryotoxicity testing of IVF disposables: how do manufacturers test? Hum Reprod. 2020;35(2):283–92.

26. Ozil JP, Huneau D. Activation of rabbit oocytes: The impact of the Ca2+ signal regime on development. Development. 2001;128(6):917–28.

27. Ozil JP, Markoulaki S, Toth S, Matson S, Banrezes B, Knott JG, et al. Egg activation events are regulated by the duration of a sustained [Ca2+]cyt signal in the mouse. Dev Biol. 2005;282(1):39–54.

28. Tóth S, Huneau D, Banrezes B, Ozil JP. Egg activation is the result of calcium signal summation in the mouse. Reproduction. 2006;131(1):27–34.

29. Lawitts JA, Biggers JD. Culture of preimplantation embryos. Methods Enzymol. 1993;225:153–64.

30. Ozil JP, Swann K. Stimulation of repetitive calcium transients in mouse eggs. J Physiol. 1995;483(Pt 2):331–46.

31. Ducibella T, Schultz R, Ozil J. Role of calcium signals in early development. Semin Cell Dev Biol. 2006;17(2):324–32.

32. Barker DJ. The fetal and infant origins of adult disease. BMJ. 1990;301(6761):1111.

33. Saunders CM, Larman MG, Parrington J, Cox LJ, Royse J, Blayney LM, et al. PLCζ: a sperm-specific trigger of Ca2+ oscillations in eggs and embryo development. Development. 2002;129(15):3533–44.

34. Cuthbertson KSR, Cobbold PH. Phorbol ester and sperm activate mouse oocytes by inducing sustained oscillations in cell Ca2+. Nature. 1985;316(6028):541–2.

35. Swann K. Sperm-Induced Ca2+ Release in Mammalian Eggs: The Roles of PLCζ, InsP3, and ATP. Cells. 2023;12(24):2809.

36. Bernhardt ML, Stein P, Carvacho I, Krapp C, Ardestani G, Mehregan A, et al. TRPM7 and CaV 3.2 channels mediate Ca2+ influx required for egg activation at fertilization. Proc Natl Acad Sci U S A. 2018;115(44):E10370–8.

37. Carvacho I, Ardestani G, Lee HC, McGarvey K, Fissore RA, Lykke-Hartmann K. TRPM7-like channels are functionally expressed in oocytes and modulate post-fertilization embryo development in mouse. Sci Rep. 2016;6(1):34236.

38. Savy V, Stein P, Delker D, Estermann MA, Papas BN, Xu Z, et al. Calcium signals shape metabolic control of H3K27ac and H3K18la to regulate EGA. bioRxiv. 2025 Mar 16. doi:10.1101/2025.03.14.643362.

